# Value and spatial preferences guide habitual reaching and manual object selection in primates

**DOI:** 10.64898/2026.06.20.733501

**Authors:** Yun-Hyuk Kim, Jun Park, Yu Gyeong Kim, Youngjeon Lee, Hyoung F. Kim

**Affiliations:** School of Biological Sciences, Seoul National University, Seoul 08826, Republic of Korea; National Primate Research Center, Korea Research Institute of Bioscience and Biotechnology (KRIBB), Cheongju, Republic of Korea; Department of Functional Genomics, KRIBB School of Bioscience, Korea University of Science and Technology (UST), Daejeon, Republic of Korea

## Abstract

Habits are automatic actions shaped by prior experience, enabling efficient and fast responses in stable environments. While habitual gaze rapidly directs the eyes toward valuable objects, the mechanisms underlying habitual manual choice for reaching and grasping remain unclear. Here we show that macaque monkeys develop habitual manual choices toward previously rewarded objects through multi-day object–value learning, driven by learned value and spatial preferences. This reaching habit persisted without reward and showed shorter latencies than in value-deliberative tasks, consistent with automatic control. Regression analyses further revealed that manual choice was guided by learned object values, whereas visual salience had no effect, unlike gaze habits. Instead, intrinsic spatial preferences continued to bias reaching behavior even after long-term value learning. These findings demonstrate that habitual manual choice arises from the integration of spatial preferences and long-term value memory, defining a distinct mechanism of automatic behavior beyond habitual gaze.

## Introduction

Through repeated experience, actions become automatic, forming habits that enable fast and efficient behavior in stable environments ^1–7^. For example, when a thirsty individual repeatedly reaches for a nearby glass of water, the action gradually becomes automatic—executed rapidly and without deliberation. This automatic choice allows faster and more efficient selection of valuable objects than competitors in stable contexts.

Previous studies have mainly focused on habits formed through extensive repetition, where repeated movements render an action automatic ^8–12^. Yet repetition alone may not fully explain why the thirsty individual so readily reaches for the glass. In this example, the motivational value of water – not mere repetition of the reaching movement – drives the action. This distinction highlights an alternative pathway for habit formation: value associations may trigger automatic behaviors independently of motor repetition ^13–15^.

Value-based habits have been documented in the primate oculomotor system ^7,16–23^. Once visual objects are associated with reward, macaques and humans show persistent gaze biases toward high-valued objects, even when value is no longer relevant and feedback is withheld ^7,16,19,21,24^. This behavior, which persists without reinforcement and occurs with shorter latencies than goal-directed saccades, has been taken as a hallmark of automatic control in visual habit^19,23^. Functionally, such a system provides a fast, low-cost way to orient toward valuable objects without deliberation, thereby enhancing efficiency and survival ^3,7,25,26^. Experimentally, visual habit is typically studied by training primates across multiple days on object–reward associations and then testing them in reward-neutral sessions. Animals continue to direct gaze toward previously high-valued objects, demonstrating stable, automatic biases ^16,19,21,23^. These findings have been central to identifying basal ganglia mechanisms underlying visual habits.

However, in natural conditions, valuable objects cannot be obtained by gaze alone; they must be physically reached for and grasped. This raises a critical question: if visual habit can emerge from value-based learning, might manual choice behavior also become habitual through similar mechanisms? Despite this, little is known about whether the value-based mechanisms that drive habitual gaze extend to manual choice. In everyday life, acquiring reward depends not on looking but on reaching and grasping, making the hand the effector most directly tied to obtaining value. Yet prior studies of hand habits emphasized motor repetition, leaving unanswered whether long-term value learning alone can induce automatic hand actions ^8–10^.

At the circuit level, value-based visual habits are thought to arise from long-term value signals encoded in the striatum and conveyed through the substantia nigra pars reticulata (SNr) to control saccades. Anatomical evidence indicates that the same striatal value signals also project through the globus pallidus internal segment (GPi) to hand motor pathways ^19,27–29^, providing a plausible route by which value memories could shape habitual manual choice. However, eye and hand effectors differ in their functional demands, raising the possibility that value-based habits may exhibit effector-specific characteristics ^30^. Since both SNr and GPi receive striatal value signals, we hypothesized that long-term learning would induce habitual manual choice, with effector-specific divergence emerging at the output level.

To address this, we tested whether long-term object-value learning can induce habitual manual choice behavior in primates and whether the underlying mechanisms differ from those driving visual habits. We developed a multi-phase behavioral paradigm in which monkeys learned visual object–reward associations through manual choices across multiple days and were subsequently tested in a value-neutral manual choice task without reward or feedback. This allowed us to assess whether previously learned value alone could drive persistent, automatic hand movements, and whether such behavior reflected long-term value memory by retesting after a one-week retention period. As a hallmark of habit is automaticity – fast, efficient responses that bypass deliberative evaluation ^3,4,19,21^ – we further compared response latencies in the habitual task with those in a value-deliberative reversal learning task. Finally, to identify the factors shaping habitual manual choices, we performed regression analyses with object value, spatial preference, object preference, and visual salience as predictors.

## Results

### Formation of long-term value memory for visual objects via manual choice

To investigate whether habitual manual choice behavior developed through stable object-value associations, we implemented a structured task schedule consisting of Pre-learning, Learning, and Retention phases (Fig. 1A). During the Pre-learning phase, we established a behavioral baseline reflecting the monkeys’ innate manual choice preferences using a habitual choice task conducted prior to value learning. In the Learning phase, monkeys learned the reward values associated with visual fractal objects, forming the basis of value-based habitual choice behavior. On the first day of this phase (Day 0), monkeys performed an object-value learning task in which particular visual objects were paired with specific reward outcomes (Fig. 1A). On subsequent days (Day 1-9), the habitual choice task was conducted before the object-value learning task to assess the development of manual choice behavior based on long-term value memory, independent of any immediate reward feedback. This daily sequence was repeated for nine consecutive days. One day after the final value learning session (Day 10), we tested whether the monkeys showed habitual choice behavior. To examine the persistence of the learned habitual behavior, monkeys underwent a one-week retention period without exposure to the trained objects, after which they were again tested with the habitual choice task.

**Fig. 1.**
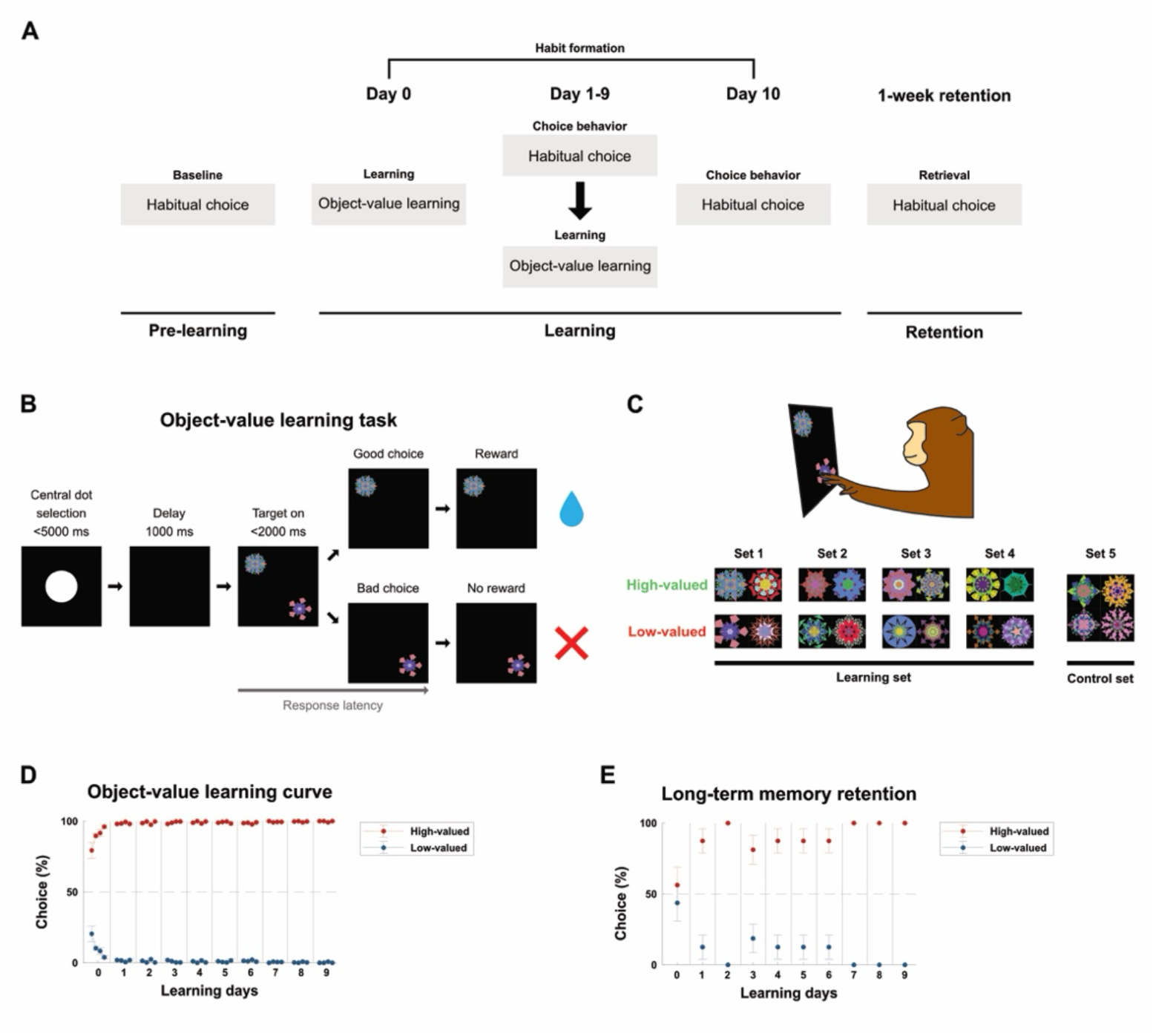
Experimental design and long-term object–value learning in macaque monkeys. **a,** Task schedule. Two monkeys learned object values through a Pre-learning session followed by 10 days of Learning sessions, and these values were later tested in a 1-week retention phase. The habitual choice task was conducted before object-value learning to assess monkeys’ innate object and spatial preferences. During the Learning phase, the habitual choice task was conducted before the object–value learning task each day to examine how habitual manual choice behavior developed. The habitual choice task was conducted again 1 week after the last testing session. **b,** Object–value learning task. Each trial began with touching a central dot followed by a delay, after which two fractal objects (high- and low-valued) appeared at pseudorandom locations. Selecting the high-valued object delivered a reward, whereas selecting the low-valued objects yielded none. **c,** Stimuli sets. Each learning set contained four fractals (two high- and two low-valued) that were learned across 10 days. An additional control set of four fractals (Set 5) was not used for object-value learning. **d,** Object-value learning curve. Choice of high-valued object increased rapidly and reached a plateau (three-way ANOVA with value, day, and bin as factors, \*\*\**p* = 3.43 × 10⁻⁷⁶ for value × day, \*\*\**p* = 3.06 × 10⁻⁵ for value × bin, \*\*\**p* = 4.67 × 10⁻¹⁸ for value × day × bin). Each bin corresponds to a single behavioral session; four sessions (bins) were conducted per day for each object set. Choice proportions were computed separately for each session and then averaged across sessions and monkeys. Data are shown as mean ± SEM across sessions. **e,** Initial trial bias. Initial trial choices on each learning day shifted toward high-valued objects from Day 1, and were sustained thereafter, confirming long-term value memory (post hoc Bonferroni pairwise comparison, \*\*\**p* = 2.694 × 10⁻^3^ between Day 0 and 1, *p* = 0.683 for Day 1 and 2; one-sample t-test).

In the object-value learning task (Fig. 1B), monkeys performed a two-choice discrimination paradigm using fractal images associated with different reward values. Each learning set consisted of four fractals: two high-valued and two low-valued (Fig. 1C). In each trial, a pair of fractal objects was presented simultaneously at pseudorandomly assigned locations among 4 possible screen positions, and monkeys were required to select one by touching it. Choosing a high-valued object resulted in a water reward, whereas selecting a low-valued object yielded no reward. This consistent reward structure enabled the monkeys to acquire stable value representations of individual visual objects.

Monkeys rapidly acquired object-value associations, with the proportion of high-valued object choices increasing sharply on the first day of learning and quickly reaching a plateau (Figs. 1D and S1A). A three-way ANOVA with value, day, and bin as factors revealed a significant main effect of value (F(1, 560) = 169,328.54, p < 0.001), as well as significant interactions of value × day (F(9, 560) = 59.80, p = 3.43 × 10⁻⁷⁶), value × bin (F(3, 560) = 8.02, p = 3.06 × 10⁻⁵), and value × day × bin (F(27, 560) = 5.97, p = 4.67 × 10⁻¹⁸). Together, these results indicate that monkeys learned the object-value associations over time, as reflected in their progressively stronger preference for high-valued objects across days and within sessions.

Furthermore, we obtained evidence of choice behavior based on long-term value memory by examining the monkeys’ first choices in the initial trial of each learning session, as this was the first time they encountered the object each day. On the first day of learning, monkeys did not exhibit a significant preference between high- and low-valued objects in the initial trial (Day 0 in Fig. 1E). However, from the second day onward, a significant value bias was observed, with monkeys increasingly selecting high-valued objects for the first choice (Figs. 1E and S1B). A post hoc Bonferroni pairwise comparison confirmed a significant increase in high-valued object selection between Day 0 and 1 (p=2.694 × 10⁻^3^), while no significant difference was found between Day 1 and 2 (p=0.683), suggesting that the value bias rapidly stabilized after the second day. This pattern indicates that object-value associations were consolidated across sessions and retained over time. These findings confirm that the monkeys acquired stable value representations of fractal objects through the learning task, forming the basis for examining whether these value memories subsequently guide habitual manual choice behavior under more naturalistic condition.

### Long-term value memory influences habitual manual choice behavior

To examine habitual manual choice behavior guided by long-term value memory, we used a habitual choice task in which the four previously learned fractals were presented at pseudorandomly selected positions among 4 of 8 possible touchscreen locations (Fig. 2A and Movie S1). Monkeys erased each object by touching it, with trial progress indicated by a central gauge bar that filled incrementally with each object removal. Once all four objects were erased, the monkey obtained a liquid reward by touching the fully filled gauge bar (Figs. 2A and C). Importantly, no reward was provided during the object choices themselves, and neither object identity nor the sequence of selections influenced value reinforcement, thereby eliminating direct feedback for individual choices. Thus, the optimal strategy was to select all four objects as quickly and automatically as possible. Because habitual behaviors persist in the absence of reward feedback, this paradigm allowed us to assess automatic, habitual choice behavior driven by long-term value memory.

**Fig. 2.**
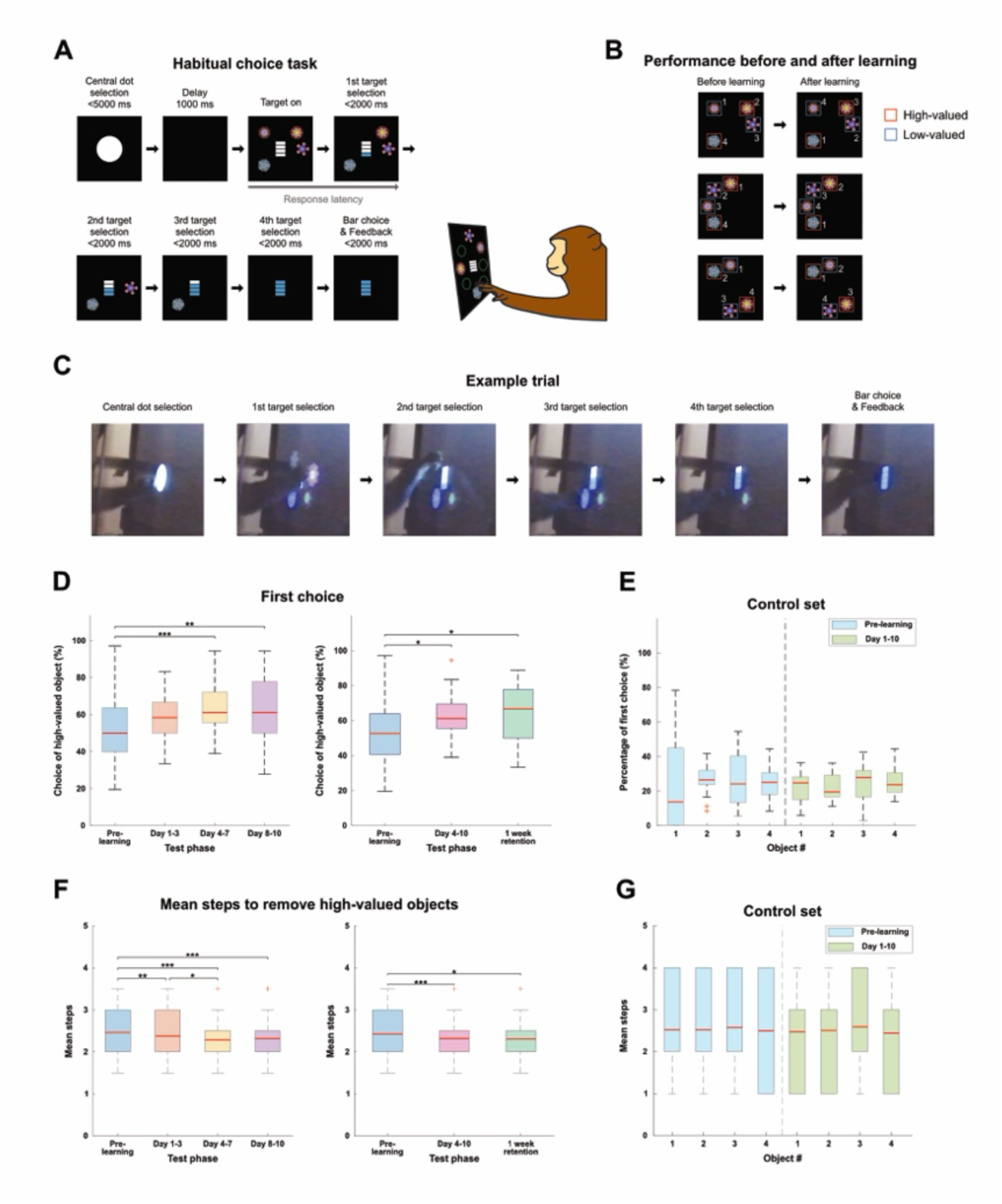
Habitual manual choice guided by long-term object-value memory. **a,** Habitual choice task. Each trial began with a central dot touch followed by a delay, after which four fractal objects from a learned set were presented at pseudorandom positions. Monkeys sequentially removed all objects by touching; after the last object was removed, touching the central gauge bar delivered a reward. No feedback was given during object choices, and reward was provided at the end of each successful trial independent of object identity or order. **b,** Example of performance before and after learning. Each stimuli configuration was tested both before and after object-value learning. **c,** Example trial. An example trial showing a monkey touching each visual fractal object with its right hand. **d,** Percentage of first choice of high-valued object. Proportion of trials in which a high-valued object was chosen first across Pre-learning, Day 1–3, Day 4–7, and Day 8–10 phases (left panel; n=80, post hoc Bonferroni pairwise comparison, \*\**p* <0.01, \*\*\**p* <0.001), and across Pre-learning, Day 4-10, and the 1-week retention phases (right panel; n=56, post hoc Bonferroni pairwise comparison, \**p*<0.05). Boxplots show medians (red line), interquartile ranges (boxes; 1st-3rd quartile), and whiskers extending to the most extreme non-outlier values (‘+’). **e,** First choice distribution for control set. Percentage of first choices by object identity (Objects #1–4) for the control set during Pre-learning and Day 1–10 phases (n=20, one-way ANOVA, *p* = 9.44 × 10^−1^). Boxplot format as in panel (d). **f,** Mean steps to remove high-valued objects. Average steps required to remove all high-valued objects across Pre-learning, Day 1–3, Day 4–7, and Day 8–10 phases (left panel; n=2880, post hoc Bonferroni pairwise comparison, \**p* <0.05, \*\**p* <0.01, \*\*\**p* <0.001), and across Pre-learning, Day 4-10, and the 1-week retention phases (right panel; n=1440, post hoc Bonferroni pairwise comparison, \**p* <0.05, \*\*\**p* <0.001). Boxplot format as in panel (d). **g,** Mean step distribution for control set. Mean selection step by object identity (Objects #1–4) for the control set during Pre-learning and Day 1–10 phases (n=720, one-way ANOVA, *p* = 5.64 × 10^−2^). Boxplot format as in panel (d).

We first examined the first choice of each trial to assess whether monkeys exhibited a bias toward high-valued objects, reasoning that if their choices were guided automatically by long-term value memory, they would tend to select previously learned high-valued objects before low-valued ones (Fig. 2B). Examination of test phases using the habitual choice task showed that the bias to select high-valued objects emerged following an initial 3-day learning period (Figs. 2D and S2A-B, left panel – Day 1-3), consistent with previous reports that primates require several days to acquire value-based habitual behavior ^20^. Notably, after 3 days of learning in the habitual choice task, monkeys exhibited a significantly higher selection of previously learned high-valued objects on their initial choice compared to baseline object selections prior to learning (Fig. 2D, left panel – Day 4-7) (one-way ANOVA, p < 0.001 for Pre-learning vs. Day 4-7). This preference persisted throughout the Learning phase (Fig. 2D, left panel – Day 8-10) (one-way ANOVA, p < 0.001 for Pre-learning vs. Day 8-10). These findings indicate that monkeys formed habitual preferences for previously learned high-valued objects, independent of immediate reward feedback.

To evaluate the persistence of this habitual choice behavior, monkeys were re-tested after a one-week retention period without additional learning and object exposure. The proportion of first choice toward high-valued objects remained comparable to that observed during the Learning phase, even one week after the last learning (Fig. 2D, right panel) (one-way ANOVA, p = 3.40 × 10^−2^ for Pre-learning vs. Day 4-10; p = 3.31 × 10^−2^ for Pre-learning vs. 1-week retention). This indicates that habitual choice bias, driven by long-term value memory, was sustained over the one-week period.

To rule out performance fluctuations or task familiarity as alternative explanations, we included a control object set that had not been associated with rewards. This control set was tested on the same days as the learning object sets (Day 1-10). Monkeys did not exhibit any significant first-choice bias to specific objects across sessions, confirming that the observed habitual choice behavior was specific to the learned value associations (Figs. 2E and S2C-D).

We further assessed the degree of habitual prioritization of high-valued objects by quantifying the number of steps needed to select and remove all high-valued objects. This number significantly decreased following object-value learning compared to baseline (Figs. 2F and S2E-F). This reduction emerged during the three-day learning period and stabilized in subsequent phases, supporting the notion that value-based habitual behavior became gradually established over time (Fig. 2F, left panel) (one-way ANOVA, p = 3.55 × 10^−3^ for Pre-learning vs. Day 1-3; p < 0.001 for Pre-learning vs. Day 4-7; p < 0.001 for Pre-learning vs. Day 8-10). Importantly, the reduction persisted after the one-week retention period, providing further evidence that the behavior was driven by long-term memory (Fig. 2F, right panel) (one-way ANOVA, p < 0.001 for Pre-learning vs. Day 4-10; p = 1.15 × 10^−2^ for Pre-learning vs. 1-week retention). The absence of such change in the control set (Figs. 2G and S2G-H) confirms that the observed effect was specifically driven by learned value associations, rather than by general task familiarity or familiarity with the objects themselves. The manual choice behavior observed here, shaped by prior experience and independent of ongoing reward, is consistent with the definition of habitual behavior.

### Response latency indicates automaticity of habitual manual choice behavior

Automatic, habitual behavior is generally characterized by fast actions executed through simple sensory-to-action circuits, in contrast to more deliberative behaviors that rely on cognitive flexibility and involve more complex brain circuits ^3,4,7,27,29^. Thus, it is important not only to examine whether actions based on previously learned values persist in the absence of reward feedback, but also to assess how quickly these learned actions are executed.

To differentiate the speed of habitual responses from deliberative goal-directed actions, monkeys were tested in a value-reversal version of the object-value learning task, which required flexible, short-term value-based decision-making (Reversal value-based manual choice task) (Fig. 3A). In each trial of the reversal task, the two fractal objects were presented simultaneously at 2 of 4 possible positions, and the monkeys selected one by touching the display. Choosing the high-valued object resulted in a liquid reward, whereas selecting the low-valued object yielded no reward. Within a single session, object-value associations were reversed twice, requiring cognitive flexibility to dynamically update object-value association (Fig. 3A). We introduced this reversal component to establish a behavioral context that demands flexible value updating, providing a contrast to the automatic response observed in the habitual choice task.

**Fig. 3.**
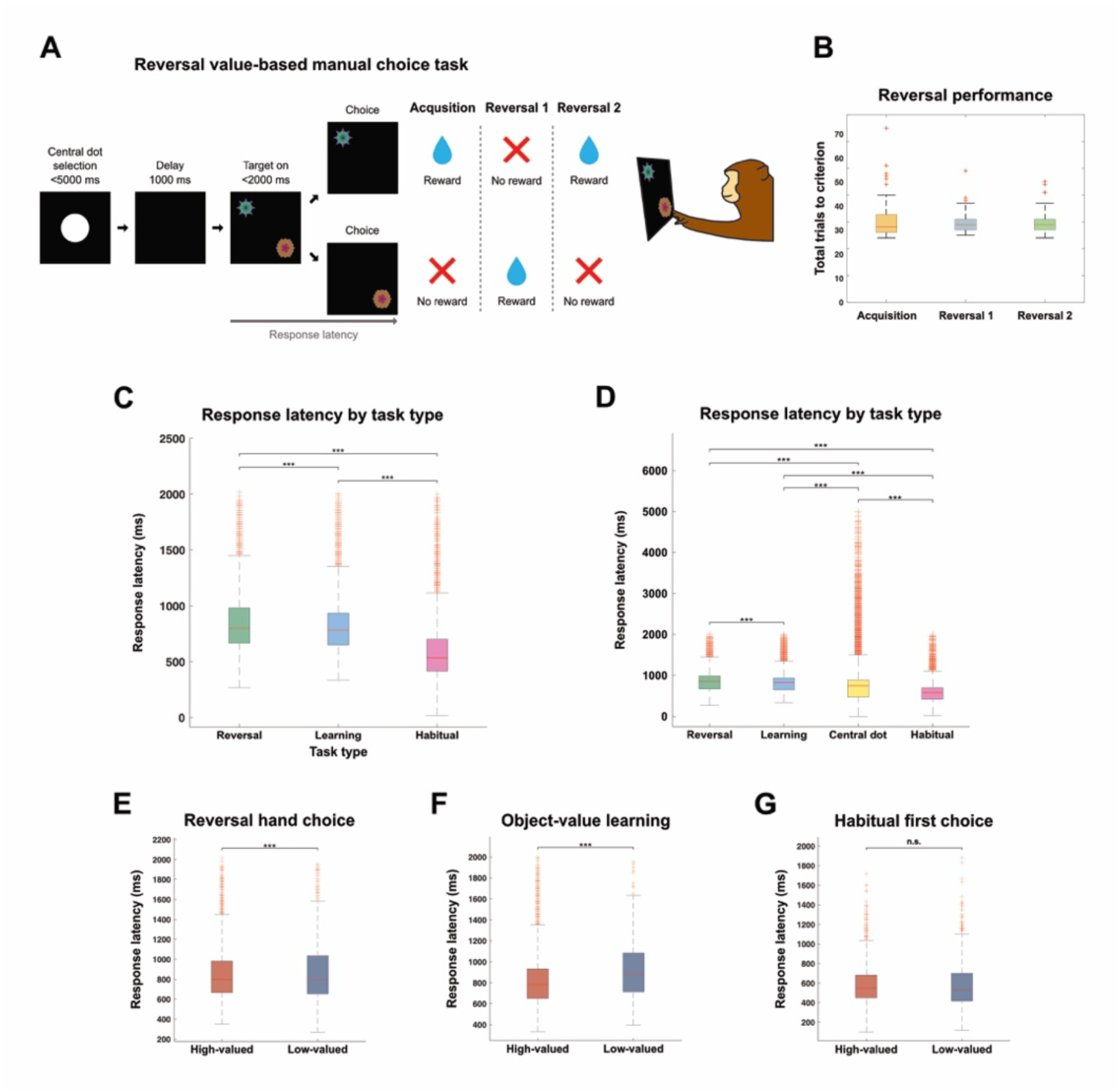
Response latencies indicate automaticity of habitual manual choice. **a,** Reversal value-based manual choice task. Monkeys initiated trials with central dot touch, followed by a delay, after which two fractal objects (high- and low-valued) were presented at pseudorandom locations. Selecting the high-valued object delivered a reward, whereas the low-valued object provided none. Object–value contingencies were reversed twice per session, requiring flexible updating. **b,** Reversal task performance. The number of trials required to reach reversal criterion (24 consecutive correct choices) did not differ significantly between acquisition and reversal blocks. Boxplots show medians (red line), interquartile ranges (boxes; 1st-3rd quartile), and whiskers extending to the most extreme non-outlier values (‘+’). Data represent mean ± SEM. **c,** Response latencies across tasks. Response latencies were significantly shorter in the habitual choice task compared to object–value learning and reversal tasks (n=45060, post hoc Bonferroni pairwise comparison, \*\*\**p* <0.001). Boxplot format as in panel (b). **d,** Response latencies comparison with central dot touch. Central-dot responses were faster than object selections in object-value learning and reversal tasks, but habitual object selections were faster than central-dot responses (post hoc Bonferroni pairwise comparison, \*\*\**p* <0.001). Boxplot format as in (b). **e, f,** Response latencies to high- and low-valued objects during reversal value and object value learning. Responses were faster for high-valued than low-valued objects for both reversal (e) and object-value learning (f) task, indicating ongoing value deliberation during task response (paired-sample t-test, \*\*\**p* <0.001). Boxplot format as in panel (b). **g,** Comparison of response latencies to high- and low-valued objects in the habitual choice task. No response latency difference was observed between high- and low-valued first choices, indicating non-deliberative responding during the habitual task (paired-sample t-test, n.s., not significant). Boxplot format as in panel (b).

Progression to the next reversal block required the monkeys to select the high-valued object in 24 consecutive trials. The number of trials required to reach this reversal criterion did not significantly differ across blocks in either monkey (Figs. 3B and S3A), indicating successful acquisition of flexibly changing object-value association.

We defined response latency as the time from object presentation to object selection and compared latencies across three behavioral tasks: the object-value learning task, the reversal value-based manual choice task, and the habitual choice task (Figs. 1B, 2A, and 3A). Because the object-value learning and reversal tasks required deliberative discrimination of associated value, we hypothesized that monkeys would exhibit longer response latencies in these tasks compared to the habitual task, which does not require explicit value evaluation ^21,31^. Consistent with this prediction, monkeys exhibited significantly shorter response latencies during the habitual choice task (741.15 ± 3.52 ms) compared to both the object-value learning task (749.57 ± 3.27 ms) and the reversal task (760.11 ± 4.84 ms), indicating a more automated behavioral response (one-way ANOVA, F(2, 44,478) = 6075.57, p < 0.001) (Figs. 3C and S3B-C).

To further evaluate response automaticity, we compared latencies for touching the central dot across tasks (Fig. 3D). Since the central dot touch is an instructed, non-value–dependent action, it serves as a motor baseline for each task, against which object-touch latencies can be interpreted. As expected, in the object–value learning and reversal tasks, central dot latencies were shorter than object-touch latencies (one-way ANOVA, p < 0.001 for Central dot vs. Learning; p < 0.001 for Central dot vs. Reversal) (Fig. 3D). In contrast, in the habitual choice task, object-touch latencies were even shorter than central dot touch latency (one-way ANOVA, p < 0.001 for Central dot vs. Habitual choice) (Fig. 3D), indicating that object-directed actions in the habitual choice task were executed with highly automatic timing.

Finally, we examined whether value influenced response latency within each task. In both the object-value learning and reversal tasks, response latencies were significantly faster for high-valued objects compared to low-valued objects (Learning: 819.00 ± 1.87 ms for high-valued objects; 931.36 ± 13.33 ms for low-valued objects; paired t-test, t(281) = −11.40, p = 5.19 × 10^−25^) (Reversal: 849.20 ± 3.28 ms for high-valued objects; 872.19 ± 6.31 ms for low-valued objects; paired t-test, t(1237) = −13.72, p = 5.46 × 10^−40^) (Figs. 3E-F). These value-dependent differences in response latency suggest that choices in these tasks were guided by deliberative evaluation of object value. In contrast, no significant latency difference was observed between high- and low-valued object selections in the habitual choice task (582.04 ± 3.80 ms for high-valued objects; 587.67 ± 4.49 ms for low-valued objects; paired t-test, t(1910) = 1.67, p = 0.09,), suggesting that such conscious value-based deliberation did not take place (Fig. 3G). This absence of value-dependent latency modulation further supports the conclusion that behavior in the habitual task was guided by automatic processes rather than deliberative value evaluation.

### Value and spatial preference shape value-based habitual manual choice behavior

What are the key factors that drive habitual manual choice movements? Do such actions depend on the visual salience of the target, the motivational value of the object, or internal biases such as spatial or object preference? While habitual visual behavior is often guided by visual salience and object-based value associations ^7^, it remains unclear whether these same drivers apply to hand-based habits. Given that gaze and hand systems serve distinct computational roles—gaze being object-centered and visually guided, and hand movements being spatially constrained and biomechanically shaped ^30^ — we tested whether the behavioral drivers of value-based habitual manual choices differ from those of habitual gaze behavior.

To identify the key contributors to manual choice behavior, we performed trial-level regression analyses using visual salience, innate object preference, innate spatial preference, and value as predictors (Fig. 4A). Visual salience was computed using a graph-based visual saliency (GBVS) algorithm ^32^, with each object assigned a ranked saliency score per trial. Innate spatial preference was assessed through a pre-learning movement test, where objects were presented in all possible position configurations (4 objects ⅹ 8 positions, 1680 configurations). The selection order for each position was used to compute preference scores for the eight spatial locations (Fig. 4B). Notably, spatial preferences differed between subjects: monkey B exhibited a leftward bias, while monkey M showed a rightward bias (Fig. 4B), suggesting that spatial preference is individually variable and innate. Similarly, object preference was calculated from pre-learning tests based on the selection order of each object (higher object preference score for objects selected earlier during pre-learning movement tests) (Fig. S4A). Value was coded as a binary variable, with high-valued objects coded as 1 and low-valued objects as 0 (Fig. 4A). Regression results revealed distinct patterns across test phases. Visual salience did not significantly influence choice behavior in any phase (95% CI: −0.19, 0.10; −0.14, 0.25; −0.59, 0.19 for Pre-learning, Learning, and Retention, respectively) (Fig. 4C). This finding contrasts with previous studies reporting a strong influence of visual salience on habitual gaze behavior, suggesting distinct mechanisms underlying hand-based habits ^7,33^.

**Fig. 4.**
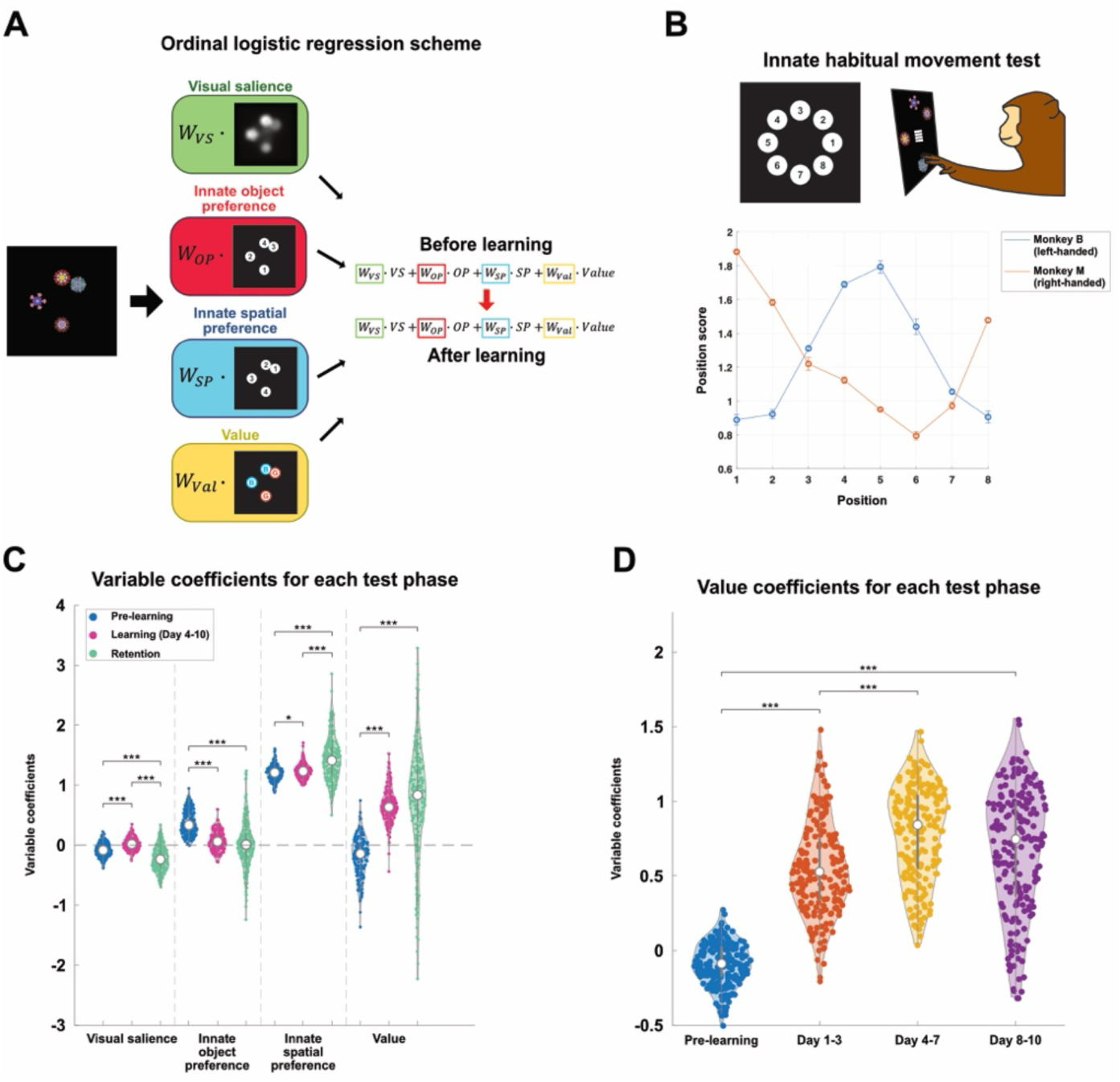
Value and spatial preferences drive habitual manual choice. **a,** Regression scheme. Ordinal logistic regression model predicting object selection order (1st–4th) from four predictors: visual salience (VS), innate object preference (OP), innate spatial preference (SP), and object value (Val). **b,** Innate spatial preference for each monkey. Pre-learning tests of the habitual choice task yielded position scores across eight screen locations (top) for each monkey, revealing subject-specific spatial biases during object selection (bottom). Monkey B was left-handed, whereas monkey M was right-handed. Data represent mean ± SEM. **c,** Regression coefficients across test phases. Regression coefficients (n=200 iterations) for VS, OP, SP, and Val during the Pre-learning, Learning (Day 4–10), and Retention phases. SP was a consistent predictor across all phases; Val became significant after learning and persisted through retention; OP was predictive only before learning; and VS showed no effect (post hoc Bonferroni pairwise comparison, \**p* <0.05, \*\*\**p* <0.001). Violin plots show coefficient distributions, with thick bars indicating the interquartile range (1st–3rd quartiles), white circles the median, and colored points the coefficient from each iteration. Coefficient panels show mean estimates ± 95% CI. **d,** Value coefficients across learning phases. Value coefficients across finer learning bins (Pre-learning, Day 1–3, Day 4–7, Day 8–10), demonstrated a progressive increase during the Learning phase (post hoc Bonferroni pairwise comparison, \*\*\**p* <0.001). Violin plot format as in panel (c).

Interestingly, innate spatial preference was the most consistent and robust predictor across all phases, indicating that spatial biases exert a strong and persistent influence on habitual manual choice movements (95% CI: 1.24, 1.53; 1.01, 1.49; 0.82, 2.19 for Pre-learning, Learning, and Retention, respectively) (Fig. 4C). This effect was further supported by response latency analyses, which revealed that positions with stronger spatial preference were associated with shorter latency in both the Pre-learning and Learning phases (Fig. S4B). In contrast, innate object preference significantly predicted behavior before learning (95% CI: 0.14, 0.51) but lost predictive power after learning (95% CI: −0.19, 0.34; −0.80, 0.97 for Learning and Retention, respectively) (repeated measures ANOVA, p < 0.001 for Pre-learning vs. Learning; p < 0.001 for Pre-learning vs. Retention) (Fig. 4C). This suggests that object value learning suppressed object biases but not spatial bias.

Value emerged as a significant predictor following learning (Fig. 4C). In the Pre-learning phase, value did not significantly influence behavior (95% CI: −0.38, 0.16), but it became a strong predictor after learning (95% CI: 0.15, 1.30) and retained its influence during the Retention phase after the last learning session (95% CI: 0.25, 2.60) (repeated measures ANOVA, p < 0.001 for Pre-learning vs. Learning; p < 0.001 for Pre-learning vs. Retention) (Fig. 4C). Although individual variability in coefficient magnitude was observed, both monkeys displayed a significant increase in value coefficients after learning (Figs. S4C-D). Further analysis of value coefficients across learning stages revealed a marked increase after the initial 3-day learning period, consistent with behavioral transitions observed earlier (repeated measures ANOVA, p < 0.001 for Pre-learning vs. Day 4-7; p < 0.001 for Pre-learning vs. Day 8-10) (Figs. 4D and S4E-F). In the late learning period (Day 4-10), the value coefficient remained elevated relative to the early learning period (Day 1-3), indicating that once established, value-based habits persisted (Fig. 4D).

Together, these findings demonstrate that value and spatial biases, but not visual salience, shape habitual manual choice behavior—highlighting effector-specific mechanisms distinct from those observed in gaze. Notably, visual salience did not contribute to habitual manual choice behavior. These findings suggest that the mechanisms underlying habitual manual choice behavior differ fundamentally from those reported for habitual gaze behavior ^20^.

## Discussion

In this study, we demonstrated that habitual hand choice behavior in primates emerges from the combined influence of learned object values and stable spatial preferences, while visual salience exerts no measurable effect. These habits persisted even when feedback was withheld and were executed with reduced response latency, thereby meeting two canonical criteria of habitual control: independence from immediate reinforcement and automaticity of execution. By identifying value and spatial preference as the principal drivers of manual choice, our findings reveal that the determinants of manual choice habits differ from those of oculomotor habits, where both value and visual salience influence gaze. This dissociation suggests that although long-term value memories form a common substrate across effectors, their behavioral expression is shaped by effector-specific constraints—visual salience in gaze, spatial preference in hand. In this way, our study extends the framework of value-based habits from the oculomotor to the manual domain, providing a behavioral foundation for linking long-term object value memories to effector-specific expressions of habit.

### Functional significance of value-based habitual manual choice

Most prior studies of hand habits have emphasized motor repetition as the key driver of automaticity (e.g. sequential finger movement) ^34–36^, showing that actions repeated over time can be expressed without conscious control ^1,2,6,11^. Our findings demonstrate that long-term object value, distinct from motor repetition-guided habits, can also establish stable manual choice habits.

Two behavioral signatures confirmed this. First, monkeys’ choices did not depend on reward contingencies but instead reflected the previously learned values of fractal objects. Both monkeys preferentially selected previously high-valued objects on the first touch of a trial, despite the absence of feedback, and this bias persisted after one-week retention period (Figs. 2D-G). These results indicate that, unlike simple motor habits formed through repetition, such movements can become habitual through sensory–motor associations driven by long-term value memory.

Second, choices were automatic. Responses in the habitual task were faster than in value-deliberative tasks, and object-selection latencies in the habitual task were shorter even than simple central dot-touch latencies, indicating streamlined sensorimotor execution (Figs. 3C-D). Moreover, in value-deliberative tasks, choice latencies differed between high- and low-valued options in both this and previous studies, whereas in the habitual task no such latency difference was observed (Figs. 3E-G), consistent with execution that bypasses deliberative evaluation ^37,38^. These properties, well established for gaze habits ^20,23,29^, are now demonstrated for manual choice, showing that long-term value memory can drive rapid, stimulus-guided hand actions without immediate reinforcement.

The adaptive advantages of such value-driven hand habit are clear in natural settings. Foraging and tool use require fast, repeatable reaches toward objects with established utility or nutritional value. Automating these selections reduces cognitive load, stabilizes performance, and conserves energetic resources ^12^. In our task, this efficiency was reflected not only in first-choice biases but also in fewer steps to clear high-valued objects (Fig. 2F), indicating prioritized sequencing once the habit formed. Thus, value-based habitual control is not restricted to gaze; it generalizes to manual choice actions, where it likely supports economical interaction with valuable objects in familiar environments ^1,2,6^.

### Behavioral drivers of habitual manual choice versus habitual gaze

Regression analyses indicate a dissociation in the behavioral mechanisms underlying manual choice and gaze habits. In the oculomotor domain, previous studies have shown that both learned object value and visual salience strongly bias gaze behavior, even when feedback is absent ^7,39^. In contrast, in our manual choice task, visual salience exerted no significant influence at any phase. Instead, two factors proved decisive: spatial preference, which was a consistent predictor from Pre-learning through Retention, and learned object value, which emerged as a predictor only after value learning. Innate object preference measured before learning lost significance once value associations were established. Thus, the shared component across effectors is a value signal that emerges after training and supports habit expression without immediate reward; the dissociations are the absence of salience effects and the dominant role of space in the hand domain.

These results map onto the distinct computational demands of the two effector systems. For gaze, selection operates in an object-centered frame where value and visual salience jointly accelerate saccades toward previously rewarded objects ^7,40^. For the hand, in contrast, movement planning requires integration of spatial geometry and biomechanical costs such as reach distance, posture, and energetic efficiency ^30^. Consistent with this, our latency analyses showed that positions with stronger spatial preference were selected faster, indicating that motor costs and spatial ease of execution directly modulate the automaticity of habitual manual choice actions. Under natural conditions where a valuable object consistently appears in the same position (e.g. a ripe apple growing on the same branch of a tree each year), spatial preference combined with value-guided choice may allow animal to reach the object more quickly and with less energy.

By showing that value-based habits extend from gaze to manual choice movements, our study demonstrates that habits are not unitary but shaped by effector-specific constraints. Gaze habits integrate value with visual salience to optimize rapid orienting, while hand habits combine value with spatial preference to support efficient interaction with reachable objects. Linking these behavioral dissociations to neural circuit mechanisms will be essential for understanding how habits are implemented across effectors and how they support adaptive behavior in natural contexts.

Together, these findings suggest a simple principle: value is the shared driver of habit expression across effectors, but the complementary factor is effector-specific – visual salience for gaze and spatial constraints for manual choice. This principle yields testable predictions such as that manipulations that change visual salience should bias selectively habitual gaze without affecting habitual manual choice. Testing such dissociations will be essential for linking behavioral differences to their underlying circuit implementations.

### Plausible basal ganglia circuit mechanisms underlying habitual manual choice

Our results raise important questions about how value information is stored and expressed in neural circuits to guide reaching behavior. In the oculomotor system, value-based visual habits have been linked to long-term representations in the striatum, which influence activity in the SNr and downstream superior colliculus (SC) to guide saccades even in the absence of reward feedback ^14,16,21,29,41^. Likewise, habitual manual choices are likely mediated by striatal value signals, as the striatum not only processes value information for reaching behavior in the reversal value task but may also serve as the locus for long-term value memory ^37,38^. However, unlike the visual habit circuit, object value signal for manual choice may be transmitted through the GPi to thalamo-cortical and motor pathways ^28^. This circuit architecture suggests a mechanism by which long-term value memory can guide reaching behavior without the need for current reinforcement.

The reduced response latencies observed during habitual manual choice suggest that, once value associations are consolidated, their expression is routed through relatively direct sensorimotor circuit. Unlike deliberative goal-directed actions, which rely on prefrontal and associative networks in the rostral basal ganglia circuits, visual habit is mediated by caudal parts of thalamus and basal ganglia ^29^. Anatomical studies have shown that these caudal thalamus and basal ganglia structures receive inputs from visual areas, including the inferotemporal cortex, and send outputs to motor areas such as the SNr ^4,29,42–44^. Moreover, long-term value memory processed within these caudal structures enable fast, automatic gaze shifts toward valuable objects ^27^, supporting the view that caudal thalamic and basal ganglia mechanisms may also underlie habitual manual choice behavior.

While such sensorimotor circuits that encode long-term value memory for visual habit provide a plausible mechanism for habitual manual choice behavior, neural response data and causal manipulations are required to establish this. The primate behavioral paradigm developed here provides a strong foundation for such studies, enabling test of whether striatal value signals are transmitted through GPi–cortical pathways to drive habitual manual choices, and how these pathways interact with SNr-mediated circuits for visual habit.

## Materials and methods

### General procedures

Two adult monkeys (*Macaca fascicularis;* 2.2 kg female monkey B, 8 years old; 3.3 kg female monkey M, 17 years old) were used for the experiments. Animal care and experimental procedures were approved by the Korea Research Institute of Bioscience and Biotechnology (KRIBB) Institutional Animal Care and Use Committee (Approval No. KRIBB-AEC-22132), and complied with national and institutional guidelines for the care and use of laboratory animals. This study was designed and reported in accordance with the ARRIVE guidelines for reporting animal research.

### Behavioral tasks

The behavioral tasks were controlled by a MATLAB-based real-time experimentation data acquisition system (NIMH MonkeyLogic, Laboratory of Neuropsychology, National Institute of Mental Health, National Institutes of Health [LN/NIMH/NIH]). The monkey was seated in a primate cage, facing a fronto-parallel screen in a sound-attenuated and electrically shielded room. Visual stimuli were presented on an infrared touch monitor (IV-190IR, Iviewkorea, Korea). A frontal opening in the cage allowed the monkey to reach out its arm and touch the monitor. For all behavioral tasks, monkeys were trained to reach out their arm and touch the monitor to select the visual stimulus (Movie S1).

### Visual stimuli

We used fractal object images created using Fractal Geometry (https://github.com/ProfKimHF) (Figs. 1C and 3A). The mean luminance was equalized across the images using the SHINE toolbox written with MATLAB (www.mapageweb.umontreal.ca/gosselif/shine). The size of each image was approximately 20°ⅹ20° in size.

### Object-value learning task procedure

Monkeys were trained to associate visual fractal objects with either high or low value in order to establish long-term value memory of fractal objects. Each trial began with the appearance of a white central dot (Fig. 1B). The monkey was required to touch the central dot within 5 seconds to initiate the trial. Following a 1 second blank delay, two fractal objects—one high-valued and one low-valued—were presented simultaneously at diagonally opposite positions on the touchscreen. The positions were pseudorandomly selected from four possible locations (right-top, right-bottom, left-top, left-bottom). Monkeys were required to choose one of the objects within 2 seconds by touching it by hand. Selection of a high-valued object resulted in immediate delivery of a liquid reward, whereas selection of a low-valued object produced no reward.

A fixed set of four fractal stimuli (two high-valued, two low-valued) was used per session, and the same set was trained for 10 consecutive days to promote long-term learning. Each session consisted of 48 trials, with 4 sessions per day for each set. Therefore, each fractal stimulus was exposed 96 times each day, with a total of 960 associative learning trials for each stimulus across the Learning phase. Across the experiment, each monkey was exposed to four sets of four fractals, yielding 16 unique learned stimuli in total (Fig. 1C).

### Habitual choice task procedure

To assess habitual responses toward previously learned objects, monkeys performed a free sequential manual choice task using fractal stimuli previously paired with either reward or no reward. Each trial began with the presentation of a central white dot, which the monkey was required to touch within 5 seconds to initiate the trial (Figs. 2A and C). A 1 second blank delay followed, after which four fractal objects from a single set were presented simultaneously at pseudorandomly chosen positions drawn from an array of eight possible screen locations spaced at 45° intervals around the center. A vertical gauge bar with four empty segments was displayed concurrently. Monkeys erased the objects by touching them sequentially in any order. Each touch caused the selected object to disappear and filled one segment of the gauge bar, providing visual feedback of task progress. Every object had to be selected within 2 seconds, and after all objects were erased, the monkey was required to touch the fully filled gauge bar within 2 seconds to receive a liquid reward. Importantly, reward delivery depended only on completion of the trial and was not contingent on the identity or order of object selection, thereby eliminating direct reinforcement of individual choices.

The task was first administered in a baseline phase, before value learning, to measure spontaneous object selection preferences. Each object set was tested across five consecutive days, with 336 trials per session, allowing full coverage of all 1,680 possible object–location configurations. After completion of value learning, monkeys were tested in the same task over ten consecutive days, with 36 trials per session. To isolate habitual behavior from immediate reward feedback, the habitual choice task was always conducted prior to the daily object–value learning task. Each monkey completed the task with four learned object sets (16 fractals total) and one control set of novel fractals that had not been paired with reward.

To assess memory retention, monkeys were tested one week after the final training session using the same object–value learning task. During this retention test, no exposure to the trained fractals occurred in the intervening week, and performance was measured to evaluate the stability of the learned object–value associations.

### Reversal manual choice task procedure

This task was designed to assess flexible manual choice behavior driven by short-term, flexible value memory. The overall trial structure was identical to the object–value learning task, except that object–value contingencies were reversed across blocks within each session (Fig. 3A).

Each session consisted of three consecutive blocks: an initial Acquisition block followed by two reversals. At the start of the Acquisition block, two fractal objects were introduced, with one designated as high-valued and the other as low-valued. Monkeys advanced to the next block once they selected the high-valued object on 24 consecutive trials. In each subsequent block, the reward contingencies for the two objects were reversed. Sessions ended when the criterion was met in the final reversal block.

Each monkey performed the task with four distinct object pairs (eight fractals in total) (Fig. S5). Each pair was tested once per day for ten consecutive days, with the initial value assignment reversed on each subsequent day to enforce continuous updating of object–value associations.

### Data analysis Behavioral analysis

To quantify habitual behavior in the habitual choice task, we analyzed the monkeys’ first object choices. For each session, trials were divided into two halves, and within each half we calculated the proportion of trials in which a high-valued object was selected first. This analysis was performed separately for each learning set. For the control set, we computed the proportion of first choices for each fractal object across trials.

To quantify selection order in the habitual choice task, we calculated the mean step at which high-valued objects were chosen within each trial. For learning sets, the mean step was computed per trial and averaged across sessions. For the control set, the mean selection step for each fractal was computed across trials.

Response latencies for all behavioral tasks (object-value learning, reversal, and habitual manual choice task) were defined as the interval between stimulus onset and the first touch. Specifically, in the object–value learning and reversal tasks, response latency was measured as the time from fractal presentation to selection. In the habitual choice task, response latency was defined as the time from fractal presentation to the first fractal touch.

Statistical comparisons for first-choice proportions, mean selection steps, and response latencies were conducted using one-way ANOVA. Post hoc pairwise comparisons were performed with Bonferroni correction, and statistical significance was set at p < 0.05.

### Ordinal logistic regression analysis

To identify factors influencing the order of object selection after learning, we performed ordinal logistic regression using the Proportional Odds Logistic Regression (*polr*) function from the MASS package in R. Separate models were fit for each day and each subject, across four learned object sets and one control set, by the following model:

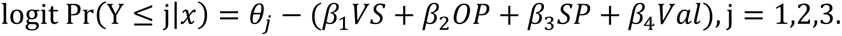

The dependent variable was the ordinal rank of touches within a trial (1st–4th), treated as an ordered categorical outcome. Four predictors were included: visual salience (VS), object preference (OP), spatial preference (SP), and object value (Val) (Fig. 4A).

Visual salience scores were computed using the Graph-Based Visual Saliency (GBVS) algorithm ^32^. For each trial, salience values were calculated for the four displayed fractals, ranked from 1 (least salient) to 4 (most salient), and entered as an ordinal predictor.

Spatial preference was derived from the Pre-learning baseline phase, in which monkeys completed trials covering all possible object–location configurations (4 objects × 8 positions, 1680 trials). Selection order across these trials was used to assign preference scores to each location. For regression, the four positions presented on a given trial were ranked (1–4) according to this preference profile and entered as the predictor for spatial preference.

Object preference was calculated in the same way as spatial preference but based on fractal identity rather than location. These scores were ranked and entered as a separate predictor.

Object value was entered as a binary categorical factor, coded 1 for high-valued and 0 for low-valued objects.

To ensure stability of coefficient estimates, 25 trials were randomly sampled without replacement from each session, and this sampling was repeated 100 times. Coefficients were averaged across iterations, and runs with incomplete data or insufficient rows were excluded. Statistical comparisons were conducted using repeated-measures ANOVA, and post hoc pairwise comparisons were performed with Bonferroni correction, with statistical significance set at p < 0.05.

### Animals and sample size justification

Two adult monkeys were used in this study. This sample size is consistent with established practices in non-human primate cognitive neuroscience, where highly trained animals perform a large number of repeated trials, enabling robust within-subject statistical analyses.

In accordance with the ARRIVE guidelines and the principle of reduction, the number of animals was kept to the minimum necessary to achieve the scientific objectives while ensuring reproducibility. Statistical analyses were performed across repeated trials and sessions within each animal, rather than treating individual animals as the primary unit of analysis.

## Data availability

Due to ethical and regulatory restrictions, the dataset supporting this study cannot be publicly archived. Access to the data requires prior approval from the Institutional Animal Care and Use Committee of Seoul National University, Korea Research Institute of Bioscience and Biotechnology (KRIBB), and National Research Foundation of Korea. Requests will be reviewed within several months and, if approved, data will be made available for academic, non-commercial use. Source data are provided with this paper.

## Code availability

Custom code associated with this study is available at https://github.com/ProfKimHF.

## Acknowledgements

This work was supported by the Basic Science Research Program (RS-2024-00339355), ASTRA Program (RS-2024-00436783), Bio&Medical Technology Development Program (RS-2025-02263832), Global-LAMP Program (RS-2023-00301976) through the National Research Foundation (NRF) of Korea, National Research Foundation of Korea (NRF) grant funded by the Korea government (MSIT) (RS-2024-00460364), and the Korea Research Institute of Bioscience and Biotechnology Research Initiative Program (KGM4562532).

The authors declare no competing financial interests.

## Author Contributions

H.F.K. and Y. Lee supervised the entire project. J. Park and Y. G. Kim performed the behavior experiment. YH. Kim and H.F.K. analyzed the data and prepared the figures. YH. Kim and H.F.K wrote the first draft, and YH. Kim, Y. G. Kim, Y. Lee, and H.F.K. interpreted data and wrote the final manuscript.

## Supplemental information

**Supplementary Fig. 1.**
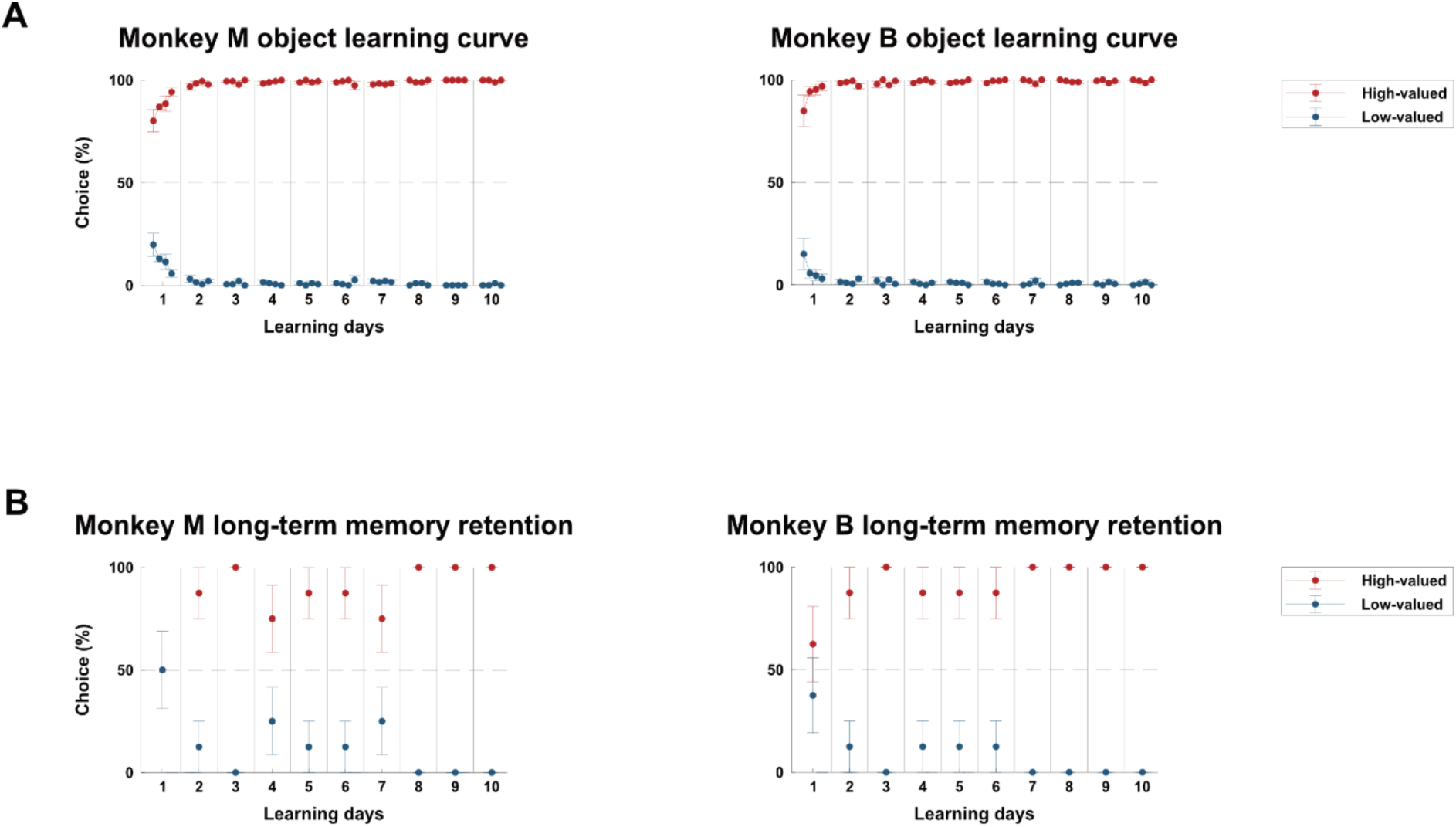
Individual monkey data for object–value learning and retention. **a,** Object-value learning curves. Proportion of high-versus low-valued object choices during the 10-day Learning phase for each monkey. Choices of high-valued object increased rapidly and reached a plateau for both monkeys. Error bars indicate ± SEM. **b,** Long-term memory retention performance. Initial trial choices shifted toward high-valued objects from Day 1 and were sustained across subsequent days, confirming long-term value memory in both monkeys.

**Supplementary Fig. 2.**
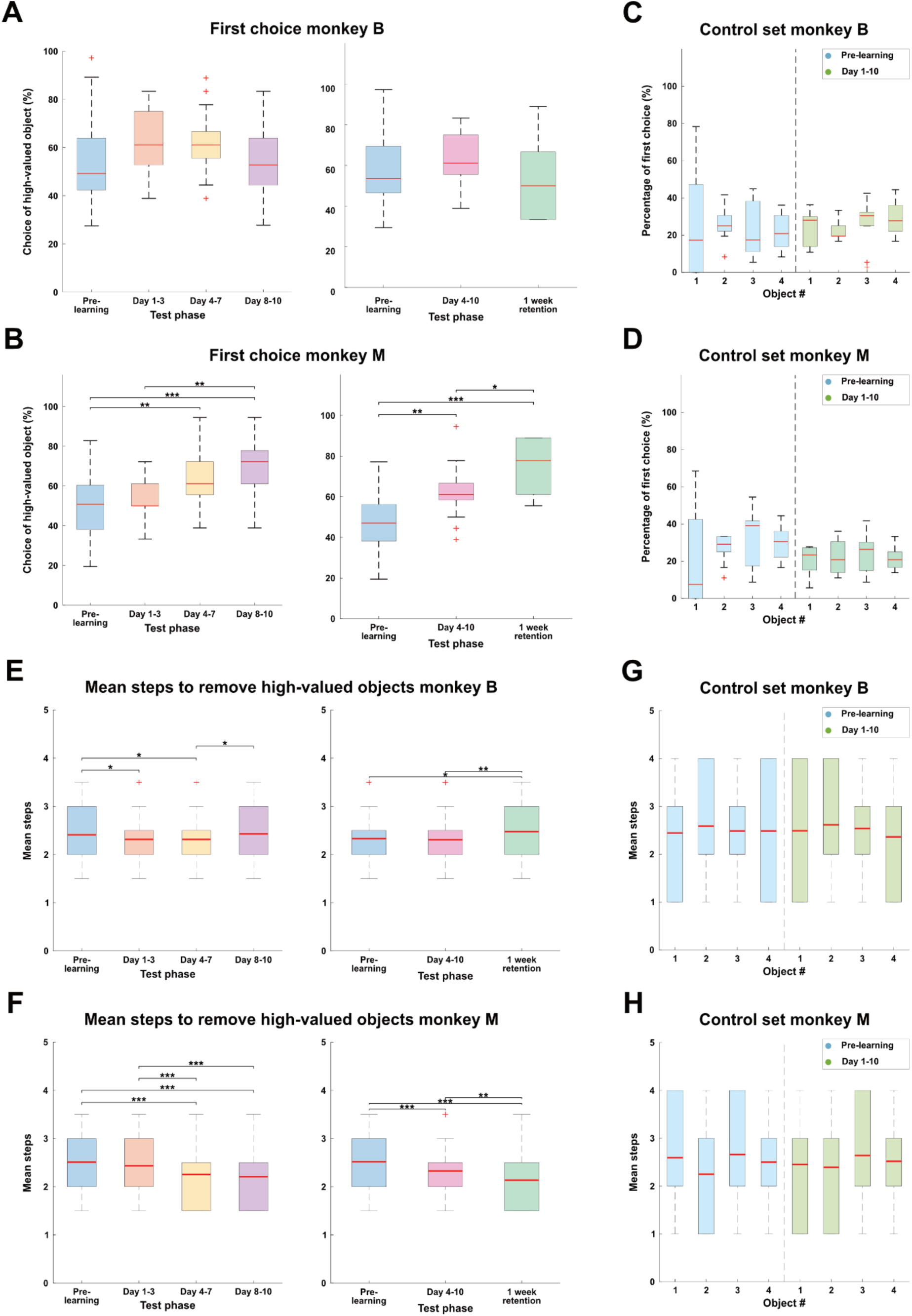
Individual monkey data for habitual manual choice behavior. **a, b,** Percentage of first choice of high-valued object. Proportion of trials in which a high-valued object was chosen first across Pre-learning, Day 1–3, Day 4–7, and Day 8–10 phases (left panel; n=40, post hoc Bonferroni pairwise comparison, \*\**p* <0.01, \*\*\**p* <0.001), and across Pre-learning, Day 4-10, and the 1-week retention phases (right panel; n=28, post hoc Bonferroni pairwise comparison, \**p*<0.05, \*\**p* <0.01, \*\*\**p* <0.001) for monkey B (a) and monkey M (b). Boxplots show medians (red line), interquartile ranges (boxes; 1st-3rd quartile), and whiskers extending to the most extreme non-outlier values (‘+’). Data represent mean ± SEM. **c, d,** First-choice distribution for control set. Percentage of first choices by object identity (Objects #1–4) for the control set during Pre-learning and Day 1–10 phases for monkey B (c) (n=10, one-way ANOVA, *p* = 0.93) and monkey M (d) (n=10, one-way ANOVA, *p* = 0.61). Boxplot format as in panel (a). **e, f,** Mean steps to remove high-valued objects. Average steps required to remove all high-valued objects across Pre-learning, Day 1–3, Day 4–7, and Day 8–10 phases (left panel; n=1440, post hoc Bonferroni pairwise comparison, \**p* <0.05, \*\*\**p* <0.001), and across Pre-learning, Day 4-10, and the 1-week retention phases (right panel; n= 720, post hoc Bonferroni pairwise comparison, \**p* <0.05, \*\**p* <0.01, \*\*\**p* <0.001) for monkey B (e) and monkey M (f). Boxplot format as in panel (a). **g, h,** Mean step distribution for control set. Mean selection step by object identity (Objects #1–4) for the control set during Pre-learning and Day 1–10 phases for monkey B (g) (n=360, one-way ANOVA, *p* = 7.38 × 10^−2^) and monkey M (h) (n=360, one-way ANOVA, *p* = 5.99 × 10^−2^). Boxplot format as in panel (a).

**Supplementary Fig. 3.**
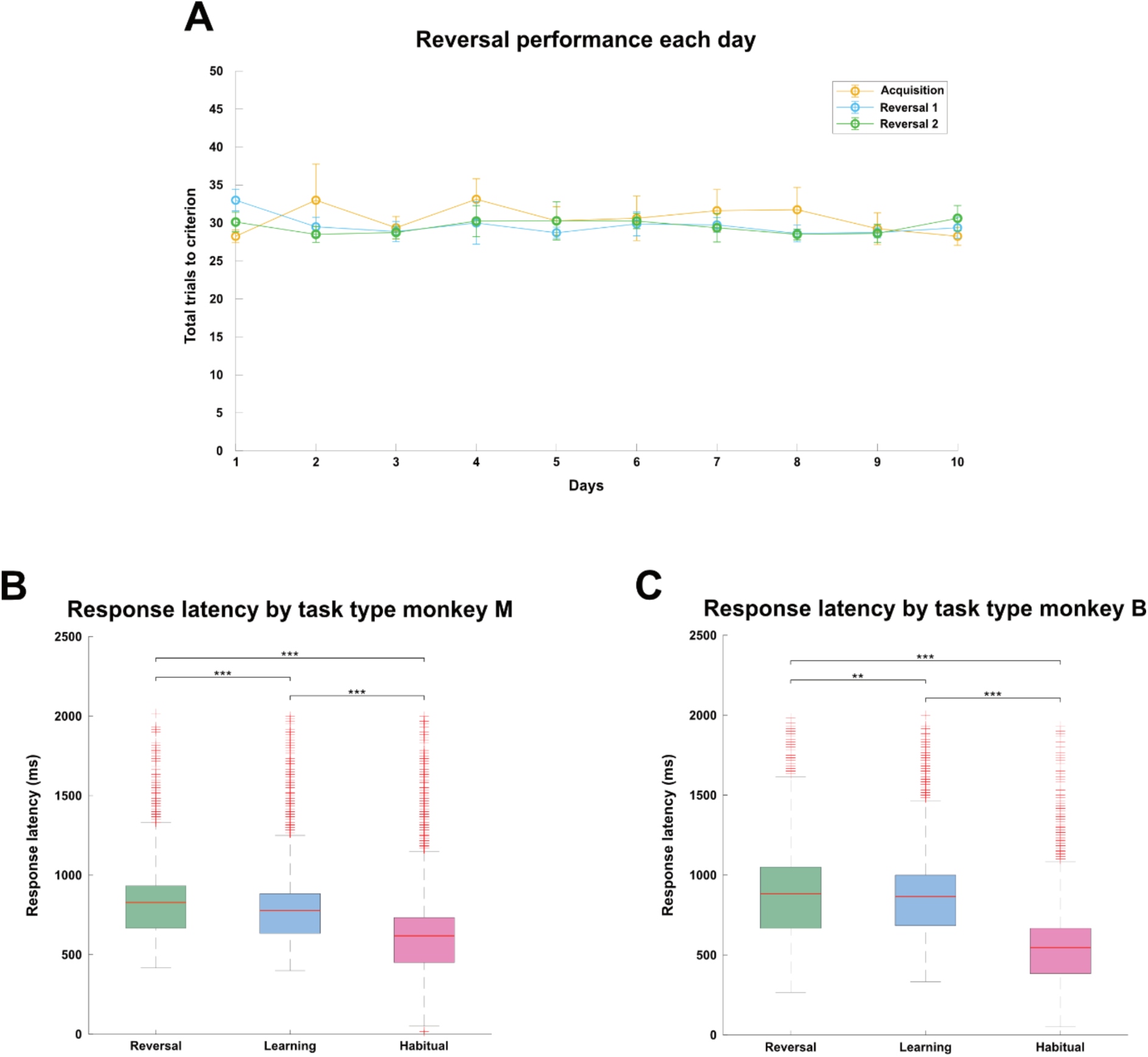
Monkey performance in the reversal task and response latencies. **a,** Reversal learning performance. Total number of trials required to reach criterion (24 consecutive correct choices) across acquisition and reversal blocks for each training day. No significant differences were observed across training days. **b, c,** Response latencies across tasks for each monkey. Response latencies were significantly shorter in the habitual choice task compared with the object–value learning and reversal tasks for both monkey B (b) and monkey M (c) (n=22530, post hoc Bonferroni pairwise comparison, \*\**p* <0.01, \*\*\**p* <0.001). Boxplots show medians (red line), interquartile ranges (boxes; 1st-3rd quartile), and whiskers extending to the most extreme non-outlier values (‘+’). Data represent mean ± SEM.

**Supplementary Fig. 4.**
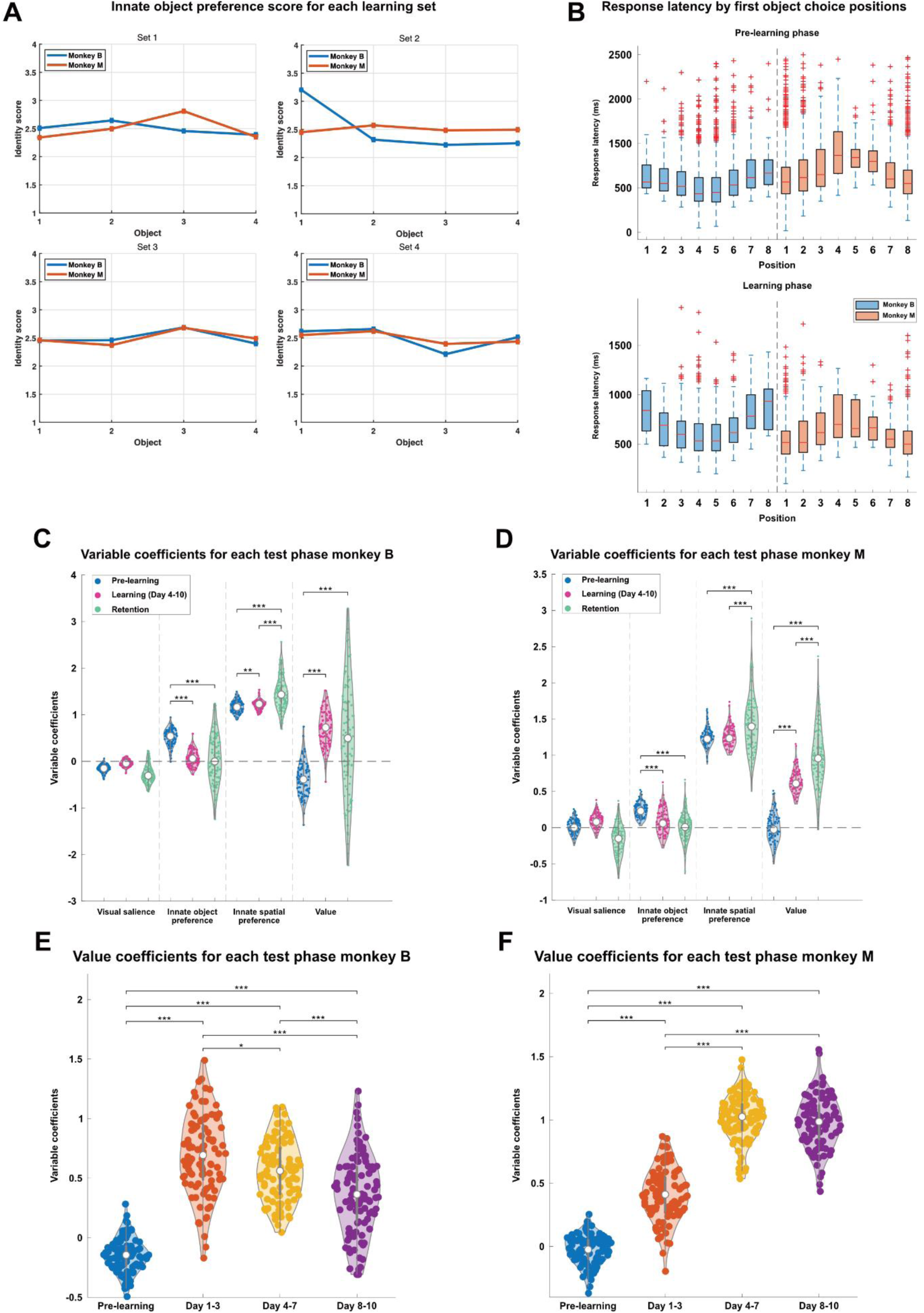
Individual regression results and spatial preference effects. **a,** Innate object preference. Innate object preference scores for each monkey, computed from Pre-learning baseline tests. **b,** Response latency by position. Response latencies for first-object choices plotted by object position for each monkey, showing faster responses at preferred positions during both the Pre-learning (top) and the Learning phase (bottom). Boxplots show medians (red line), interquartile ranges (boxes; 1st-3rd quartile), and whiskers extending to the most extreme non-outlier values (‘+’). Data represent mean ± SEM. **c, d,** Regression coefficients across test phases for each monkey. Regression coefficients (n=100 iterations) for VS, OP, SP, and Val in Pre-learning, Learning (Day 4–10), and Retention phases for monkey B (c) and monkey M (d) (post hoc Bonferroni pairwise comparison, \*\**p* <0.01, \*\*\**p* <0.001). Violin plots show coefficient distributions, with thick bars indicating the interquartile range (1st–3rd quartiles), white circles the median, and colored points the coefficient from each iteration. Coefficient panels show mean estimates ± 95% CI. **e, f,** Value coefficient across learning phases for each monkey. Value coefficients across finer learning bins (Pre-learning, Day 1–3, Day 4–7, Day 8–10) for monkey B (e) and monkey M (f) (post hoc Bonferroni pairwise comparison, \**p* <0.05, \*\*\**p* <0.001). Violin plot format as in panel (c).

**Supplementary Fig. 5.**
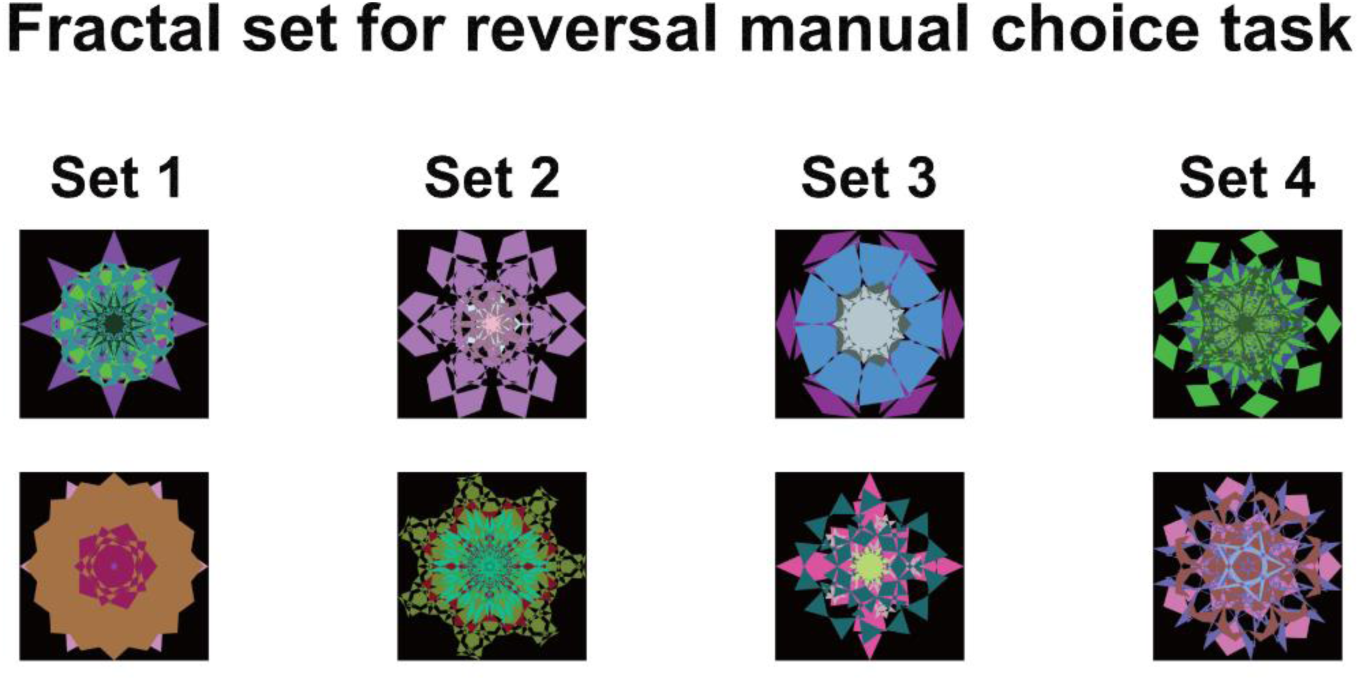
Fractal stimulus sets for the reversal manual choice task. Four distinct pairs of fractal stimuli were used in the reversal task. Within each pair, one object was initially assigned as high value and the other as low value. Object-value contingencies were reversed across blocks and across days to enforce flexible updating.

## References

1. Graybiel, A. M. Habits, rituals, and the evaluative brain. Annual Review of Neuroscience 31, 359–387 (2008).

2. Dickinson, A. Actions and Habits: The Development of Behavioural Autonomy. Trans. R. Soc. Lond. B 308, 67–78 (1985).

3. Kim, H. F. Brain substrates for automatic retrieval of value memory in the primate basal ganglia. Molecular Brain 14, 168 (2021).

4. Lee, K., An, S. Y., Park, J., Lee, S. & Kim, H. F. Anatomical and Functional Comparison of the Caudate Tail in Primates and the Tail of the Striatum in Rodents: Implications for Sensory Information Processing and Habitual Behavior. Mol Cells 46, 461–469 (2023).

5. Yin, H. H. & Knowlton, B. J. The role of the basal ganglia in habit formation. Nature Reviews Neuroscience 7, 464–476 (2006).

6. Wood, W. & Rünger, D. Psychology of habit. Annu Rev Psychol 67, 289–314 (2016).

7. Hwang, S. H., Ra, Y., Paeng, S. & Kim, H. F. Motivational salience drives habitual gazes during value memory retention and facilitates relearning of forgotten value. iScience 25, 105104 (2022).

8. Du, Y., Krakauer, J. W. & Haith, A. M. The relationship between habits and motor skills in humans. Trends in Cognitive Sciences 26, 371–387 (2022).

9. Hardwick, R. M., Forrence, A. D., Krakauer, J. W. & Haith, A. M. Time-dependent competition between goal-directed and habitual response preparation. Nature Human Behaviour 3, 1252–1262 (2019).

10. Norman, D. A. Categorization of Action Slips. Psychological Review 88, 1 (1981).

11. Smith, K. S. & Graybiel, A. M. A dual operator view of habitual behavior reflecting cortical and striatal dynamics. Neuron 79, 361–374 (2013).

12. Packard, M. G. & Knowlton, B. J. Learning and memory functions of the basal ganglia. Annual Review of Neuroscience 25, 563–593 (2002).

13. Adams, J. A. Historical review and appraisal of research on the learning, retention, and transfer of human motor skills. Psychological Bulletin 101, 41 (1987).

14. Hikosaka, O., Yamamoto, S., Yasuda, M. & Kim, H. F. Why skill matters. Trends in Cognitive Sciences 17, 434–441 (2013).

15. Gillian, C. M. et al. Disruption in the balance between goal-directed behavior and habit learning in obsessive-compulsive disorder. American Journal of Psychiatry 172, 282–290 (2015).

16. Yamamoto, S., Kim, H. F. & Hikosaka, O. Reward value-contingent changes of visual responses in the primate caudate tail associated with a visuomotor skill. Journal of Neuroscience 33, 11227–11238 (2013).

17. Peck, C. J., Jangraw, D. C., Suzuki, M., Efem, R. & Gottlieb, J. Reward modulates attention independently of action value in posterior parietal cortex. Journal of Neuroscience 29, 11182–11191 (2009).

18. Sato, M. & Hikosaka, O. Role of Primate Substantia Nigra Pars Reticulata in Reward-Oriented Saccadic Eye Movement. The Journal of Neuroscience 22, 2363–2373 (2002).

19. Hikosaka, O., Takikawa, Y. & Kawagoe, R. Role of the Basal Ganglia in the Control of Purposive Saccadic Eye Movements. Physiol Rev 80, 953–978 (2000).

20. Kang, J. et al. Primate ventral striatum maintains neural representations of the value of previously rewarded objects for habitual seeking. Nat Commun 12, 2100 (2021).

21. Kim, H. F. & Hikosaka, O. Distinct Basal Ganglia Circuits Controlling Behaviors Guided by Flexible and Stable Values. Neuron 79, 1001–1010 (2013).

22. An, S. young, Hwang, S. H., Lee, K. & Kim, H. F. The primate putamen processes cognitive flexibility alongside the caudate and ventral striatum with similar speeds of updating values. Prog Neurobiol 243, 102651 (2024).

23. Kim, H. F., Ghazizadeh, A. & Hikosaka, O. Dopamine Neurons Encoding Long-Term Memory of Object Value for Habitual Behavior. Cell 163, 1165–1175 (2015).

24. Fernandez-Ruiz, J., Wang, J., Aigner, T. G. & Mishkin, M. Visual habit formation in monkeys with neurotoxic lesions of the ventrocaudal neostriatum. Proceedings of the National Academy of Sciences 98, 4196–4201 (2001).

25. Mishkin, M., Malamut, B. & Bacheyalier, J. Memories and habits: Two neural systems. Neurobiol Learn Mem 65–77 (1984).

26. Gottlieb, J. Attention, Learning, and the Value of Information. Neuron 76, 281–295 (2012).

27. Kim, H. F., Griggs, W. S. & Hikosaka, O. Long-Term Value Memory in the Primate Posterior Thalamus for Fast Automatic Action. Current Biology 30, 2901–2911 (2020).

28. Haber, S. N. Corticostriatal circuitry. Dialogues Clin Neurosci 18, 7–21 (2016).

29. Kim, H. F. Principal axis of primate basal ganglia in processing cognitive flexibility and habitual stability. Brain awaf299 (2025).

30. Johansson, R. S. & Flanagan, J. R. Coding and use of tactile signals from the fingertips in object manipulation tasks. Nature Reviews Neuroscience 10, 345–359 (2009).

31. Takikawa, Y., Kawagoe, R., Itoh, H., Nakahara, H. & Hikosaka, O. Modulation of saccadic eye movements by predicted reward outcome. Exp Brain Res 142, 284–291 (2002).

32. Harel, J., Koch, C. & Perona, P. Graph-Based Visual Saliency. Advances in neural information processing systems 19 (2006).

33. Thompson, K. G. & Bichot, N. P. A visual salience map in the primate frontal eye field. Progress in Brain Research 147, 249–262 (2005).

34. Ohbayashi, M. Inhibition of protein synthesis in M1 of monkeys disrupts performance of sequential movements guided by memory. Elife 9, e53038 (2020).

35. Choi, Y., Yunha Shin, E. & Kim, S. Spatiotemporal dissociation of fMRI activity in the caudate nucleus underlies human de novo motor skill learning. Proceedings of the National Academy of Sciences 117, 23886–23897 (2020).

36. Alexander, G. E., Delong, M. R. & Strick, P. L. Parallel Organization of Functionally Segregated Circuits Linking Basal Ganglia and Cortex, Annual review of neuroscience 9, 357–381 (1986).

37. Hwang, S. H., Park, D., Lee, J. W., Lee, S. H. & Kim, H. F. Convergent representation of values from tactile and visual inputs for efficient goal-directed behavior in the primate putamen. Nature Communications 15, 8954 (2024).

38. Hwang, S. H., Lee, J. W., Kim, S. P. & Kim, H. F. Efficient and dynamic neural geometry of value and modality encoding in the primate putamen for value-guided behavior. Nature Communications 16, 8266 (2025).

39. Anderson, B. A., Laurent, P. A. & Yantis, S. Value-driven attentional capture. Proc Natl Acad Sci U S A 108, 10367–10371 (2011).

40. Anderson, B. A. The attention habit: How reward learning shapes attentional selection. Ann N Y Acad Sci 1369, 24–39 (2016).

41. Graybiel, A. M. & Grafton, S. T. The striatum: Where skills and habits meet. Cold Spring Harb Perspect Biol 7, a021691 (2015).

42. Jiang, H. & Kim, H. F. Anatomical inputs from the sensory and value structures to the tail of the rat striatum. Front Neuroanat 12, 30 (2018).

43. Griggs, W. S. et al. Flexible and stable value coding areas in caudate head and tail receive anatomically distinct cortical and subcortical inputs. Front Neuroanat 11, 106 (2017).

44. Kim, H. F., Ghazizadeh, A. & Hikosaka, O. Separate groups of dopamine neurons innervate caudate head and tail encoding flexible and stable value memories. Front Neuroanat 8, 120 (2014).

